# Proinflammatory and autoimmunogenic gut microbiome in systemic lupus erythematosus

**DOI:** 10.1101/621995

**Authors:** Bei-di Chen, Xin-miao Jia, Jia-yue Xu, Li-dan Zhao, Jun-yi Ji, Bing-xuan Wu, Yun-yun Fei, Hua-xia Yang, Hua Chen, Xiao-xia Zuo, Hui Li, Wen-you Pan, Xiao-han Wang, Shuang Ye, Dong-geng Guo, Li Wang, Jing Li, Lin-yi Peng, Wen-jie Zheng, Wen Zhang, Feng-chun Zhang, Jian-min Zhang, Wei He, Xue-tao Cao, De-pei Liu, Jun Wang, Xuan Zhang

## Abstract

Systemic lupus erythematosus (SLE), characterized by chronic inflammation and multi-organ damage, has been suggested to associate with gut dysbiosis, but knowledge is limited from small sample size and 16s rRNA-based studies. To shed new light on the role of microbiota in SLE development, we analyzed the fecal metagenome of 117 treatment-naïve SLE patients and 115 sex- and age-matched healthy controls (HC) by deep-sequencing; in addition, 52 of the aforementioned patients have post-treatment fecal metagenome for comparison. We found significant differences in microbial composition and function between SLE and HC, revealing multiple plausible contributing bacterial species and metabolic pathways in SLE. In-depth SNP-based analysis revealed an oral-microbiome origin for two marker species, strengthening the importance of bacterial translocation in disease development. Lastly, we confirmed experimentally that peptides of SLE-enriched species mimicking autoantigens such as Sm and Fas could trigger autoimmune responses, suggesting a potential causal role of gut microbiota in SLE.

---

The pathogenesis of SLE, a prototypical autoimmune disease resulting from loss of self-tolerance and sustained autoantibody production, has not been fully elucidated yet. Besides host genetic factors^2^, environmental triggers including microbiota are also drawing great attention^3^. Our study performed deep metagenomic sequencing to analyze the fecal DNA of 117 treatment-naïve SLE patients to exclude the effect of immune-suppressants; fecal samples after treatment were also collected from 52 patients. For comparison, we collected fecal samples from 115 sex- and age-matched healthy controls (HC) without any other inflammatory, infectious or systemic disease, nor received antibiotics within three months before sample collection (Supplementary Table 1). Fecal DNA were sequenced on Illumina Hiseq platform and a minimum of 56 million reads after quality filtering and removal of host DNA were obtained for each sample, which were then analyzed by MetaPhlAn2^4^ and HUMAnN2^5^ to derive microbial composition and function. Among all 284 samples, 597 species and 512 MetaCyc pathways were obtained.

We first compared alpha-diversity of fecal microbiota across groups, including evenness indicator Shannon index based on species-level abundance. Out of all comparisons, Shannon index was significantly lower in samples from treatment-naïve SLE patients than those from HC (P=0.016; Fig.1a), in line with a previous report showing that alpha diversity decreases within many diseases^6^. However, Shannon index continued to decrease in the post-treatment fecal samples compared to pre-treatment (P=0.044; Supplementary Fig 1a), suggesting that treatment did not restore microbial diversity in gut. Intriguingly, subgroup with lupus nephritis (LN) comprised the main source of Shannon index decline (adjusted-P=0.006; Fig.1b).

**Figure 1.**
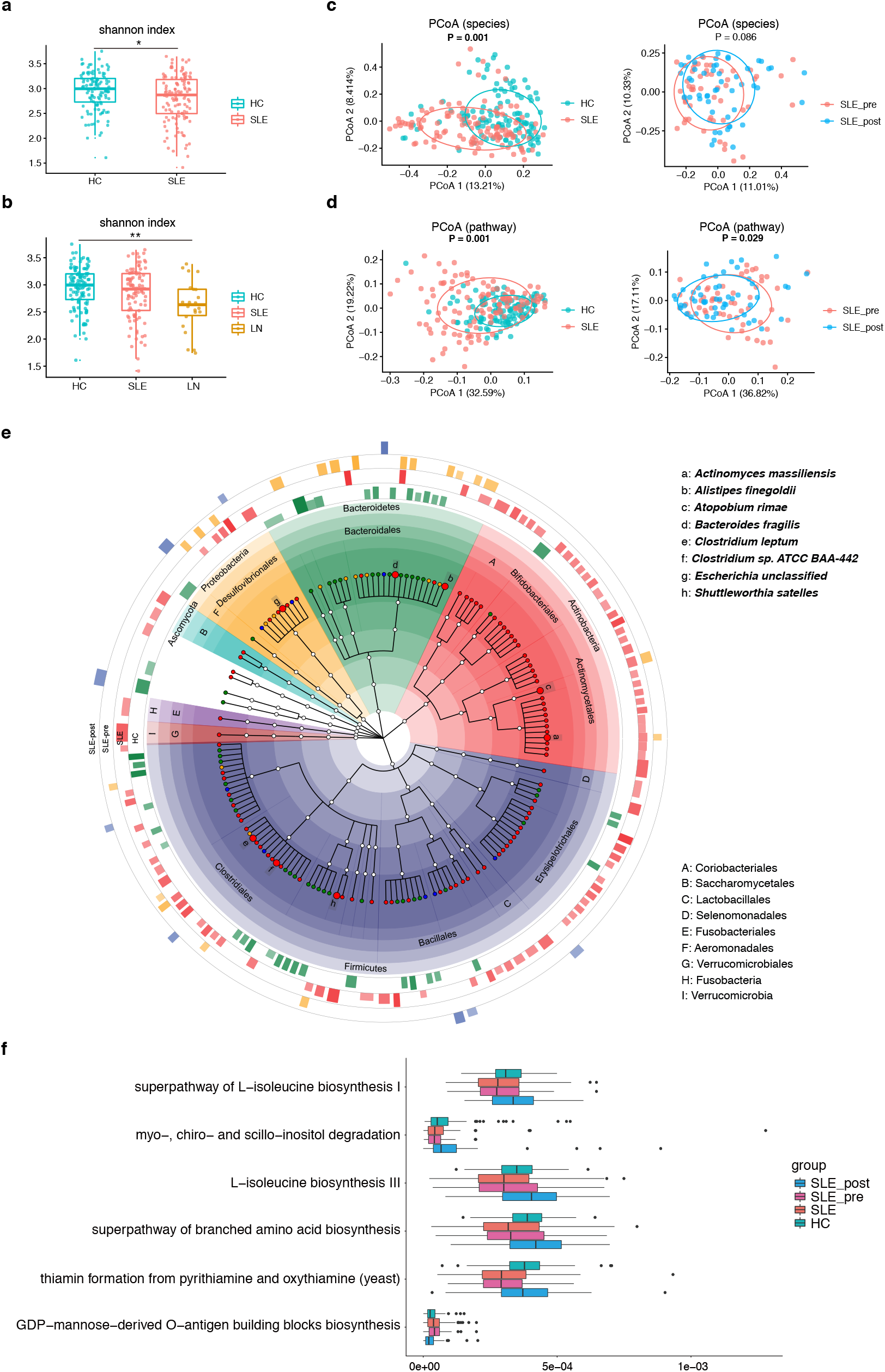
Metagenomic analyses of the fecal microbiome in systemic lupus erythematosus (SLE). **a**, Boxplot of α-diversity (Shannon index) in SLE patients and healthy controls (HC). **b**, Boxplot of α-diversity (Shannon index) in HC, SLE patients with and without lupus nephritis (LN). For **a-b**, two-tailed Wilcoxon rank-sum test and Kruskal-Wallis test were used to determine significance for two groups and multiple groups comparison, respectively. Boxes represent the interquartile ranges (IQR) and the lines inside the boxes represent the median. Each jittered point represents a fecal sample. *P < 0.05, **P < 0.01. **c,d**, β-diversity (Bray-Curtis similarity index) analyses of microbial species **(c)** and pathways **(d)** between SLE patients and HC, and between pre-treatment and post-treatment SLE patients. For **c-d**, permutational multivariate analysis of variance (PERMANOVA) was conducted with a permutation number of 999, and the significance levels are shown on plots. Each point represents a fecal sample. **e**, The taxonomical tree of all differentially enriched species. Each circular sector filled by different color represents a single phylum. The bar plots outside the central circle depict which group a specific species was enriched in and how big was the linear discriminant analysis (LDA) score. Phyla and orders were labeled on the plot. Species that increased in SLE patients and decreased after treatment were shown as large red dots. **f**, Boxplot of the pathways that were significantly different between SLE patients and HC and reversed in fecal samples after treatment. Differently enriched pathways with LDA > 2.0 were selected by LDA effect size (LEfSe) analysis. Whiskers represent the lowest or highest values within 1.5 times IQR from the first or third quartiles, and black dots represent data points beyond the whiskers. HC, n=115; SLE, n=117; pre-treatment and post-treatment SLE, n=52; SLE with LN, n=22; SLE without LN, n=95.

Beta-diversity of gut microbiota also showed distinct pattern in SLE patients. Analysis of Bray-Curtis distance based on species-level composition revealed distinct profiles between SLE and HC (*adonis*, R^2^=0.030, P=0.001; Fig 1c). The distribution of post-treatment samples demonstrated a trend to be more similar to HC group, based on principle coordinates analysis (PCoA; along the first axis PCoA1; Supplementary Fig 1b), but statistically not significant (*adonis*, R^2^=0.014, P=0.086; Fig 1c), partially due to the high within-group variability of SLE (Supplementary Fig 1b), an observation also reported by others^7^. Further analysis with SLE-related clinical parameters (Supplementary Table 2; Supplementary Fig 2a) revealed positive correlations between serum platelet and creatinine with gut microbiota variation in SLE (R^2^=0.018, P=0.045 and R^2^=0.019, P=0.037, respectively). Creatinine was also significantly correlated with gut microbiota variation of all subjects (R^2^=0.008, P=0.037; Supplementary Fig 2b), supporting the potential association between renal function and gut microenvironment in SLE development^8,9^.

To uncover the key microbial taxa that contributed to the difference between SLE and HC microbiota, linear discriminant analysis (LDA) of effect size (LEfSe) was performed using a cut-off of LDA > 2.0^10^. The correlation between relative abundance of significantly different species and clinical metadata were also calculated (Supplementary Figure 3). Seven marker species were found to be enriched in SLE patients and reduced after treatment, including *Clostridium sp. ATCC BAA-442*, *Atopobium rimae*, *Shuttleworthia satelles*, *Actinomyces massiliensis*, *Bacteroides fragilis*, *Clostridium leptum* and an unclassified *Escherichia* (Fig 1e). *Clostridium sp. ATCC BAA-442*, positively correlated with disease activity score (SLEDAI) (R^2^=0.196, P=0.034). It could metabolize Daidzein, an isoflavone previously shown to ameliorate the disease activity of lupus-prone mice by binding to estrogen receptor beta^11^, into O-demethylangolensin^12^, and thus contributed to SLE. *Bacteroides fragilis* is a common commensal bacterium composed of two subgroups, non-toxigenic (NTBF) and enterotoxigenic *Bacteroides fragilis* (ETBF). The latter produces *Bacteroides fragilis* toxin (BFT) capable of increasing colon mucosal permeability^13^, activating gut Th17 response^14^ and promoting adhesion of human T cells with vascular endothelial cell line^15^. Indeed, *bft* gene was detected in 5 samples of treatment-naïve SLE patients, and none of the HC samples carried *bft* gene (coverage>0.5 with depth>1; Supplementary Table 3). Of note, abundance of *bft* gene was positively correlated with the concentration of C3 (P=0.029, R=0.211) and C4 (P=0.045, R=0.197) in SLE serum, indicating the proinflammatory role of this bacterial group in SLE.

It is noticed that among those marker species, *Actinomyces massiliensis*, *Shuttleworthia satelles* and *Atopobium rimae* were species originally isolated from human oral cavity and were all closely related to oral inflammation^16–18^, thus their enrichment in SLE gut indicates a possible translocation from oral cavity. We carried out a strain-level analysis of those species based on single nucleotide polymorphisms (SNPs) using the oral metagenome from an independent healthy cohort^19^ and the fecal metagenome of our HC and SLE cohorts. Altogether we generated 5518, 3247 and 5429 SNPs for *Actinomyces massiliensis*, *Shuttleworthia satelles* and *Atopobium rimae*, respectively, and three phylogenetic trees were constructed (Fig 2). The strains of *Actinomyces massiliensis* in the oral controls had a closer relationship with those in SLE patients rather than those in HC (p < 0.0001; Fig 2a), and so did *Shuttleworthia satelles* (p = 0.0020; Fig 2b). For *Atopobium rimae*, SLE patients demonstrated a trend towards closer phylogeny with oral controls (p=0.0902; Fig 2c). The enrichment of potential oral species in SLE feces suggests an increased level of transmission of salivary microbes to gut, which has also been found to contribute to diseases including colorectal cancer and rheumatoid arthritis^20^.

**Figure 2.**
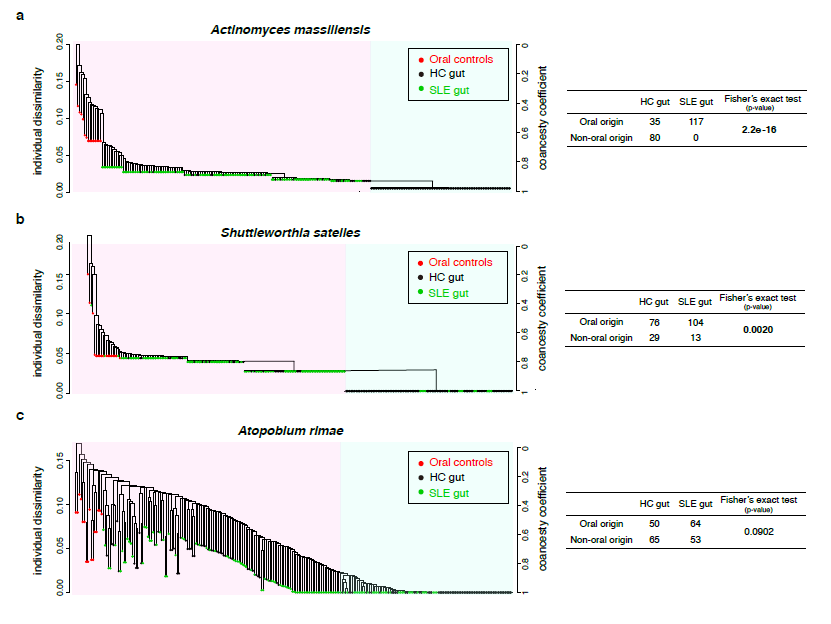
The oral origin of the enriched species in systemic lupus erythematosus (SLE) patients. **a-c**, The phylogenic trees of **(a)** *Actinomyces massiliensis*, **(b)** *Shuttleworthia satelles* and **(c)** *Atopobium rimae* based on the SNPs of each species among 16 oral samples of an independent healthy cohort, 115 fecal samples of healthy controls (HC) and 117 fecal samples of SLE patients. Each point represents a single sample and the color of the points denotes the grouping (red for oral controls, black for fecal samples of healthy controls, and green for fecal samples of SLE patients). The ancestor oral branches are shown at the left with a pink background, and the outliers are shown at the right with a blue background. For *Actinomyces massiliensis*, 117/117 of SLE gut samples and 35/115 of HC gut samples are clustered within the ancestor branch with the oral samples (p < 0.0001). For *Shuttleworthia satelles*, 104/117 of SLE gut samples and 29/115 of HC gut samples are clustered within the ancestor branch with the oral samples (p = 0.0020). For *Atopobium rimae*, 64/117 of SLE gut samples and 50/115 of HC gut samples are clustered within the ancestor branch with the oral samples (p = 0.0902). The significance levels were calculated by the Fisher’s exact test (right half).

We used HUMAnN2 to generate the gut microbial profile of functional pathways, and found that SLE patients differed substantially from HC (*adonis*, R^2^=0.047, P=0.001; Fig 1d). Functional profiles also changed significantly after treatment (*adonis*, R^2^=0.023, P=0.029; Fig 1d), indicating a higher resolution in distinguishing pre- and post-treatment gut microbiome than compositional data. LEfSe was used to analyze differentially enriched pathways (LDA>2.0; Figure 1f). “superpathway of GDP-mannose-derived O-antigen building blocks biosynthesis” was the only pathway increased in SLE patients and decreased after treatment. O-antigen comprises the outermost domain of lipopolysaccharide (LPS), the major component of the outer membrane of Gram-negative bacteria, and might act as a virulence factor aggravating SLE disease condition. The relative abundances of five pathways were lower in SLE patients and significantly reversed after treatment. Among them, three pathways were associated with branched-chain amino acids (BCAAs), including “superpathway of branched amino acid biosynthesis”, “L-isoleucine biosynthesis III” and “superpathway of L-isoleucine biosynthesis I”. BCAAs consists of valine, leucine and isoleucine, which are shown to attenuate adaptive immunity from animal-model studies^21,22^. Decreased BCAA-related pathways in gut are consistent with the reduced concentration of valine and isoleucine in SLE serum^23^, which might lead to exacerbated local and systemic inflammation.

We finally examined one key mechanism by which gut microbiota could affect SLE development, by experimentally testing the potential of microbial peptides in promoting autoimmune responses via molecular mimicry^24^. We retrieved the autoantigen peptides that were demonstrated to trigger autoimmune responses in SLE patients from the Immune Epitope Database (IEDB), including those confirmed by T-cell assays or B-cell assays, and compared them with the metagenomic sequences of species distinct between groups. With a threshold of both identity and coverage > 80%, we identified 34 peptides similar to T-cell assay epitopes and 657 peptides similar to B-cell assay epitopes (Supplementary Table 4 and 5), the latter including 16 peptides identical to confirmed auto-epitopes (Supplementary Table 6) and thus with potential immuno-pathogenicity. Using enzyme-linked immunospot (ELISPOT) assay, we demonstrated that the peptide “YLYDGRIFI” of IS66 family transposase from *Odoribacter splanchnicus*, similar with the auto-epitope “ILQDGRIFI” of human Sm B/B’ antigen^25^, was capable of increasing the secretion of IFN-γ and IL-17A from the peripheral blood mononuclear cells (PBMCs) of SLE patients (6 out of 21 and 5 out of 21 respectively; Figure 3a and b). Given that anti-Sm antibody is the most specific autoantibody found in SLE patients so far, and IFN-γ and IL-17A are implicated in the pathogenesis of SLE^26^, it is highly possible that “YLYDGRIFI” from *Odoribacter splanchnicus*, which decreased in SLE gut after treatment (Supplementary Fig 4a), can trigger SLE flares. We further discovered a peptide “DGQFCM” from *Akkermansia muciniphila*, an enriched species in SLE patients (Supplementary Fig 4b), was capable of binding to IgG produced by SLE memory B cells (P = 0.029; Figure 3c). Mimicking the extracellular part “DGQFCH” of human Fas^27^, this specific peptide might stimulate the production of anti-Fas antibody, capable of promoting apoptosis in both lupus patient lymphocytes and mouse keratinocytes^28,29^. The different responses of SLE PBMCs to the peptides above also indicate essential inter-individual variations in susceptibility towards plausible microbial auto-antigens.

**Figure 3.**
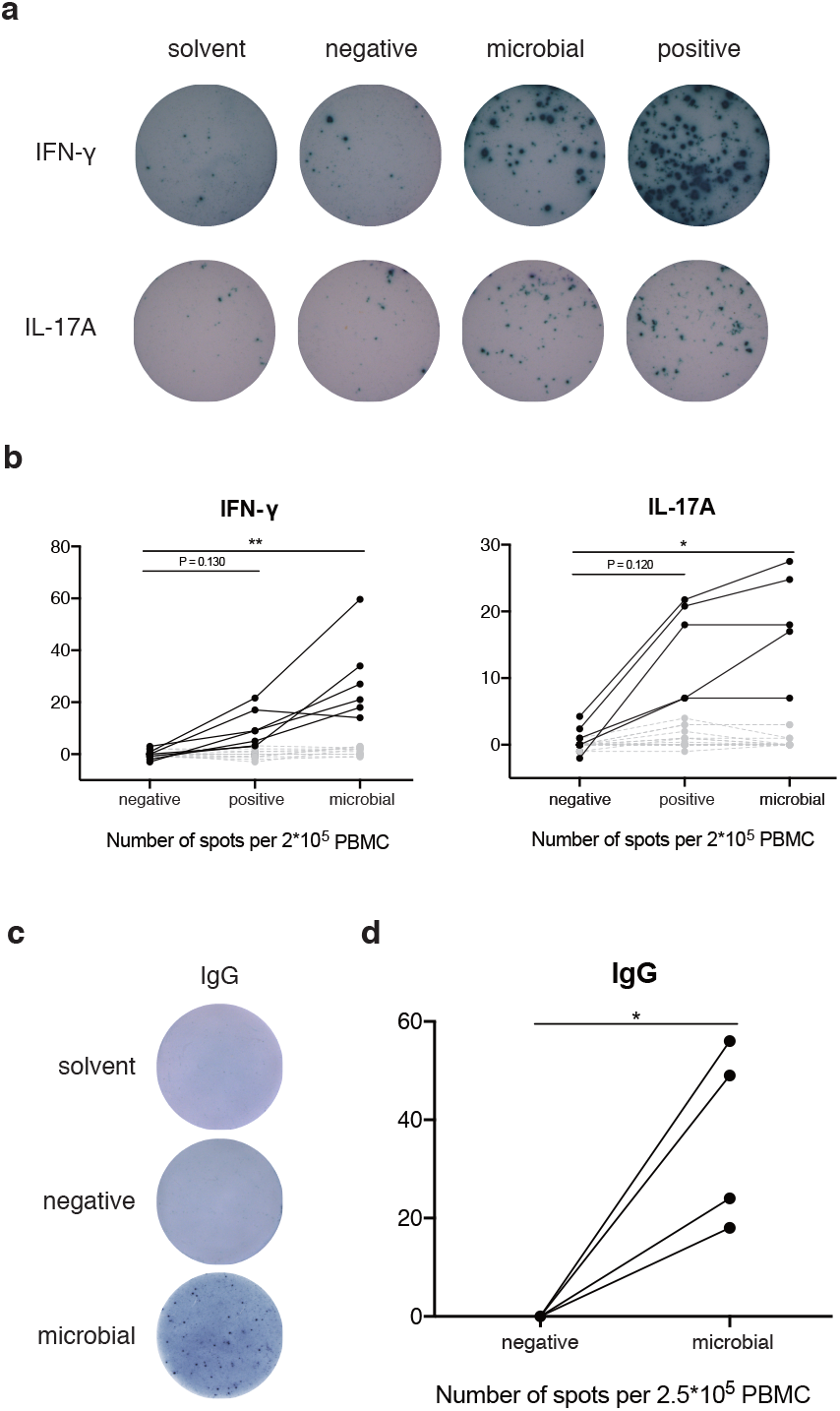
Molecular mimicry of SLE-enriched species. **a-b** Peripheral blood mononuclear cells (PBMCs) separated from SLE patients were plated in the ELISpot plates with 50 μg/ml different peptides and incubated overnight. PBMCs from 21 SLE patients were tested, and 6/21 and 5/21 were positive for IFN-γ and IL-17A secretion respectively upon stimulation by the microbial peptide "YLYDGRIFI" (similar to the Sm auto-epitope "ILQDGRIFI”) from *Odoribacter splanchnicus*. A randomly synthesized peptide "ILKEPVHGV" was set as a negative control. **a**, An example of IFN-γ and IL-17A ELISpot results. Each spot in well represents a cell that had secreted IFN-γ or IL-17A during incubation. **b**, Plots showing the number of spots inspected in different stimulation groups. **c-d** PBMCs separated from SLE patients were first pre-stimulated with a mixture of R848 at 1 μg/ml and rhIL-2 at 10 ng/ml for 96h. They were then plated in the ELISpot plates for 36h producing IgG. 6 SLE PBMCs were tested and 4 of them showed positivity for the microbial peptide “DGQFCM” from *Akkermansia muciniphila*. Detecting the IgG specific for randomly synthesized “DTYESE” was set as a negative control. **c**, An example of IgG ELISpot results. Each spot in well represents a cell that had secreted a kind of peptide-specific IgG during incubation. **d**. The plot showing the number of spots inspected in different groups. Points connected by each line represent the data from a single patient. Black points and solid lines represent the samples that were reactive to autoimmune and microbial peptides, while gray points and dash lines represent those were not reactive. The significance levels were calculated by Friedman test and paired t test for cytokine and IgG ELISpot respectively, only including the samples that were positive for the auto-epitopes. All numbers of spots were counted by subtracting the spot number of the solvent controls.

Collectively, our large-scale gut metagenomic study of the treatment-naive SLE patients with post-treatment data suggests a disturbed microenvironment in SLE gut, especially the enrichment of proinflammatory bacterial species and components, part of which being a result of translocation from oral microbiota. The multi-facet nature of SLE and inter-individual differences correspond to plural plausible microbial contributors, and the current study could not rule out the possibility that identified bacterial species/components only act secondarily by exacerbating instead of causing SLE. Nonetheless, we experimentally demonstrated that microbial peptides could trigger SLE-related cytokine production *in vitro* and bind to IgG produced by SLE memory B cells, an evidence pointing to causality between gut microbiota and SLE. Our study offers an in-depth understanding of the relationship between SLE patients and gut microbiota, and provides insights into further works elucidating the exact role of various gut microbes in autoimmune diseases.

## Acknowledgments

This study was supported by grants from the National Natural Science Foundation of China (81788101, 81630044, 81601432, 81550023, 81325019, 81771763, 81273312, 91542000, 81801633, 31771481 & 91857101), Chinese Academy of Medical Science Innovation Fund for Medical Sciences (CIFMS2016-12M-1-003, 2017-12M-1-008, 2017-I2M-3-011, 2016-12M-1-008), National Key Research and Development Program: Precise Medical Research (2016YFC0903900) National Key Science and Technology Project of China (2018YFC2000500), the Strategic Priority Research Program (Grant No. XDB29020000) and Key Research Program (Grant No. KFZD-SW-219) of Chinese Academy of Science, Grant from Medical Epigenetics Research Center, and Chinese Academy of Medical Sciences (2017PT31035).

## Authors contributions

X.Z. and J.W. designed and supervised the study; Y.F., H.Y., H.C., X.Z., L.Z, L.W., J.L., L.P., W.Z., W.Z., F.Z., H.L., W.P., X.W., D.G., and S.Y. collected the samples and performed the clinical analysis; B.C., X.J., and J.X. performed the bioinformatic analysis; B.C., J.J., and B.W. designed and performed the experiments; B.C. and J.J. analyzed and interpreted the experimental data; B.C., X.J., J.X., and J.J. drafted the manuscript; X.Z., J.W., J.Z., W.H, X.C., and D.L. revised the manuscript.

## Methods

### Study cohort

This study was approved by the Institutional Review Board of Peking Union Medical College Hospital (Ethical review number: JS-1239). Informed consent was obtained from all subjects. 117 treatment-naïve SLE patients, who had not received any steroids or immunosuppressants for the last 3 months, were recruited in Peking Union Medical College Hospital. All patients fulfilled the American College of Rheumatology revised criteria for SLE. SLE disease activity score (SLEDAI) 2000 criteria were used to evaluate disease activity. 115 sex- and age-matched healthy control were also enrolled. Subjects were excluded if they had (i) received antibiotic treatment within one month before sample collection; or (ii) current evidence of any acute or chronic inflammatory or infective diseases; or (iii) severe major systemic disease, including malignancy; or (iv) any other autoimmune disease. All participants are Han Chinese. The demographic characteristics and clinical metadata are listed in Supplementary Table 1.

### Sample collection and processing

Fecal samples were collected into a sterile stool container and frozen at –80°C within 2 hours. They were then transported frozen in dry ice and extracted at Novogene Bioinformatics Technology Co., Ltd using TIANGEN kit according to the manufacturer’s recommendations. Agarose gel electrophoresis was used to monitor DNA purity and integrity. DNA concentration was measured by Qubit® 2.0 Flurometer (Life Technologies, CA, USA) using Qubit® dsDNA Assay Kit. Samples with OD260/280 value between 1.8 to 2.0 and DNA contents above 1ug were used for library construction.

### Whole-genome shotgun sequencing

A total amount of 1μg DNA per sample was used as input material for the DNA sample preparations. Sequencing libraries were generated using NEBNext® Ultra™ DNA Library Prep Kit for Illumina (NEB, USA) following the manufacturer’s recommendations and index codes were added to attribute sequences to each sample. Briefly, the DNA sample was fragmented by sonication to a size of 350bp, then DNA fragments were end-polished, A-tailed, and ligated with the full-length adaptor for Illumina sequencing with further PCR amplification. At last, PCR products were purified (AMPure XP system) and libraries were analyzed for size distribution by Agilent2100 Bioanalyzer and quantified using real-time PCR. The clustering of the index-coded samples was performed on a cBot Cluster Generation System according to the manufacturer’s instructions. After cluster generation, the library preparations were sequenced on an Illumina HiSeq platform and paired-end reads of 250 bp nucleotide were generated.

### Metagenomic analyses

#### Quality control

More than 28 million paired raw reads were obtained for each sample. KneadData (https://bitbucket.org/biobakery/kneaddata/wiki/Home) was used for quality control of raw reads. The human (GRCh37/hg19) reference database was downloaded, and Bowtie2 contained in KneadData was used for eliminating human sequences with bowtie2-options “--very-sensitive --dovetail”. We trimmed reads with Trimmomatic in KneadData with trimmomatic-options “SLIDINGWINDOW:4:20 MINLEN:50”. To ensure comparability, 28 million paired-end reads were randomly extracted from the pre-processed data of each direction, which was 56 million reads in total for each sample.

#### Taxonomic and functional data extraction

MetaPhlAn2^1^(Metagenomic Phylogenic Analysis) was used to profile the composition of microbial communities from quality-filtered reads with `“--mpa_pkl ${mpa_dir}/db_v20/mpa_v20_m200.pkl” and “--bowtie2db ${mpa_dir}/db_v20/mpa_v20_m200”. All taxonomic data were reported as relative abundance. HUMAnN2^2^ (the HMP Unified Metabolic Analysis Network) was used for functional analysis (gene family and pathway). The gene families were annotated using UniRef90 identifiers and pathways using MetaCyc IDs. The flag “--gap-fill” was set to “on” to quantify a pathway even when a small number of its reactions were conspicuously absent. The abundance of gene family and pathway originally in reads per kilobase (RPK) was normalized to relative abundance using the humann2_renorm_table function.

#### Taxonomical and functional profiling and statistical comparison

R software version 3.5.1 was used to perform bioinformatic analyses. Alpha-diversity of each sample was calculated on the basis of species-level abundance according to the Shannon index using the vegan diversity() function Statistical significance was analyzed by non-parametric Wilcoxon rank–sum test. PCoA plots (beta-diversity) were generated by cmdscale() functions from a Bray–Curtis distance matrix of species-level taxa produced by the vegan vegdist() function. Permutational multivariate analysis of variance (PERMANOVA) was conducted using the vegan adonis() function with a permutation number of 999. Correlation between genus or species abundance and clinical parameters was calculated by Spearman correlation analysis using the WGCNA corAndPvalue () function.

LEfSe^3^ (LDA Effect Size), an algorithm for high-dimensional biomarker discovery and explanation, was used for identification of genomic features (taxa, genes, and pathways) characterizing the differences between SLE patients and HC. Both alpha value for the factorial Kruskal-Wallis test and for the Wilcoxon test between classes were set to 0.05. The threshold on the logarithmic LDA score for discriminative features was 2.0. The visualization of significant different species were generated with GraPhlAn^4^.

#### SNP-based strain-level analysis

For the analysis of the oral origin of the enriched gut species in SLE patients, 16 oral samples of an independent healthy cohort^5^, 115 fecal samples of our HC cohort and 117 fecal samples of our SLE cohort were included. StrainPhlAn was used for identifying the specific strains of species across all samples, based on the single nucleotide polymorphisms (SNPs) within the conserved and unique species marker genes. The tool is designed to track strains across large collections of samples. The raw metagenomic reads in FASTQ format were taken as input and were mapped against the set of species-specific markers (>200 per species). A variant calling approach was used by StrainPhlAn to reconstruct the sample-specific marker loci and to output the sequences of each sample-specific marker in a FASTA format. Sequences are extracted from the raw reads using a reference-free majority rule that filters out the noise regions. The resulting sequences were then concatenated and aligned by samtools. The reconstructed markers were then used to build a phylogenic tree based on the large SNP datasets by samtools and StrainPhlAn. The phylogenetic tree was built by SNPhylo with a maximum likelihood method. The number of the fecal samples that are clustered with the oral samples in each cohort is counted.

### Molecular mimicry analysis

#### Epitopes extraction

Autoantigen epitopes that were reported to trigger autoimmune reactions in SLE patients were identified from the Immune Epitope Database (IEDB, http://www.iedb.org/), including 60 epitopes proved by T-cell assays and 705 epitopes proved by B-cell assays. The amino acid sequences of these epitopes were mapped to the protein sequences of significant different species identified with LEfSe using BLAST^6^.

#### Enzyme-linked immunospot (ELISpot) assay

Heparinized blood samples were obtained from SLE participants in Peking Union Medical College Hospital. Patient peripheral blood mononuclear cells (PBMCs) were isolated by Ficoll-Hypaque density-gradient centrifugation. IFN-γ and IL-17A ELISPOT assays were performed using Human ELISpot^plus^ kit (MabTech) according to the manufacturer’s instructions. Briefly, 96-well MultiScreen Filter Plates (Millipore) pre-coated with capture antibody were washed 5× with PBS and blocked with 10% fetal bovine serum (FBS) for at least 30 min at room temperature before use. Duplicates of 2-5 × 10^5^ PBMCs per well were plated with 50 μg/ml peptides and incubated overnight (20-24 h). Plates were washed 5× with PBS and then 1 μg/ml anti-human IFN-γ mAb (7-B6-1-Biotin) or 0.5 μg/ml anti-human IL-17A mAb (MT504-Biotin) was added and incubated for 2 h. Next, each plate was washed 5× with PBS and incubated for 1 h with streptavidin–ALP, followed by 5× washes with PBS and development with 0.45-μm-filtered TMB substrate for 10-20min. The reaction was stopped by rinsing extensively with deionized water, and the plates were left to dry. Antigen-specific IgG ELISpot assays were performed using Human IgG ELISpot^BASIC^ kit (MabTech). PBMCs were first pre-stimulated with a mixture of R848 at 1 μg/ml and rhIL-2 at 10 ng/ml for 96h, promoting the ability of memory B cell to secret detectable IgG. The ELISpot plates were coated with anti-human IgG (mAbs MT91/145) overnight at 4°C. After washing and blocking with 10% FBS, duplicates of 2-5 × 10^5^ PBMCs were plated in the coated ELISpot plates for 36h. Plates were washed with PBS, then 1 μg/ml biotinylated peptides were added and incubated for 2 h. The subsequent operations were consistent with above. Plates were finally scanned and enumerated using an ImmunoSpot plate reader (Cellular Technology Limited). Each well was scored positive if the number of spot-forming cells detected were at least twice over the negative control (the randomly synthesized peptides).

### Statistical analysis

Determination of whether the SLE or HC fecal samples had a higher homology with the oral samples considering a specific species used the Fisher’s exact test. Friedman tests were utilized to calculate the significance level for ELISpot assays between different groups. The tests above were achieved using GraphPad Prism 7.0. The number of samples in each experiment is provided in the figure legends. All statistical tests in our study were performed two-sided.

### Data availability

The raw data from metagenomic sequencing are available in the SRA under BioProject ID PRJNA532888. The demographic characteristics and clinical metadata are listed in Supplementary Table 1.

