## Supplementary Fig for "Proinflammatory and autoimmunogenic gut microbiome in systemic lupus erythematosus"

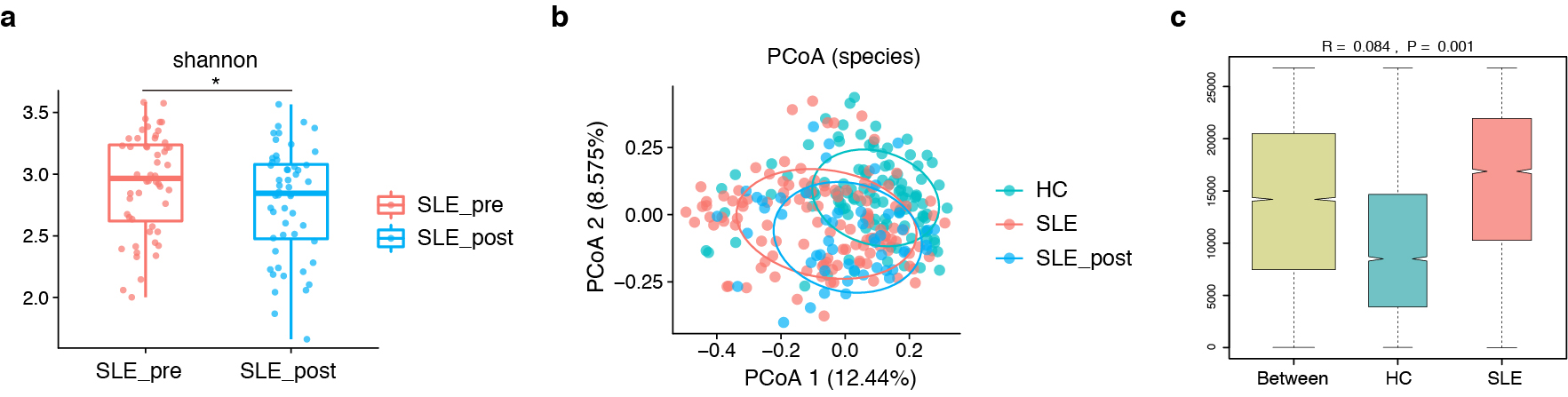

**Supplementary Figure 1. Diversity patterns of the fecal microbiome in systemic lupus erythematosus (SLE). a,** Boxplot of α-diversity (Shannon index) of pre-treatment and post-treatment fecal samples in 52 SLE patients. Two-tailed Wilcoxon rank-sum test was used to determine the significance level. Boxes represent the interquartile ranges (IQR) and the lines inside the boxes represent the median. Each jittered point represents a fecal sample. *P < 0.05. **b,** β-diversity (Bray-Curtis similarity index) analyses of microbial species between 115 healthy controls, 117 pre-treatment SLE patients and 52 post-treatment SLE patients. Permutational multivariate analysis of variance (PERMANOVA) was conducted with a permutation number of 999, and the significance levels are shown on plots. Each point represents a fecal sample. The level of confidence ellipses is 68%. **c,** Boxplot of diversity (Bray-Curtis similarity index) between and within groups of 117 SLE patients and 115 healthy controls. Analysis of similarities (ANOSIM) was conducted, with a permutation number of 999, to test whether there is a significant difference between the two groups. Boxes represent the IQR and the lines inside the boxes represent the medians, with the notches showing the 95% confidence interval for the medians. Whiskers represent the lowest or highest values.

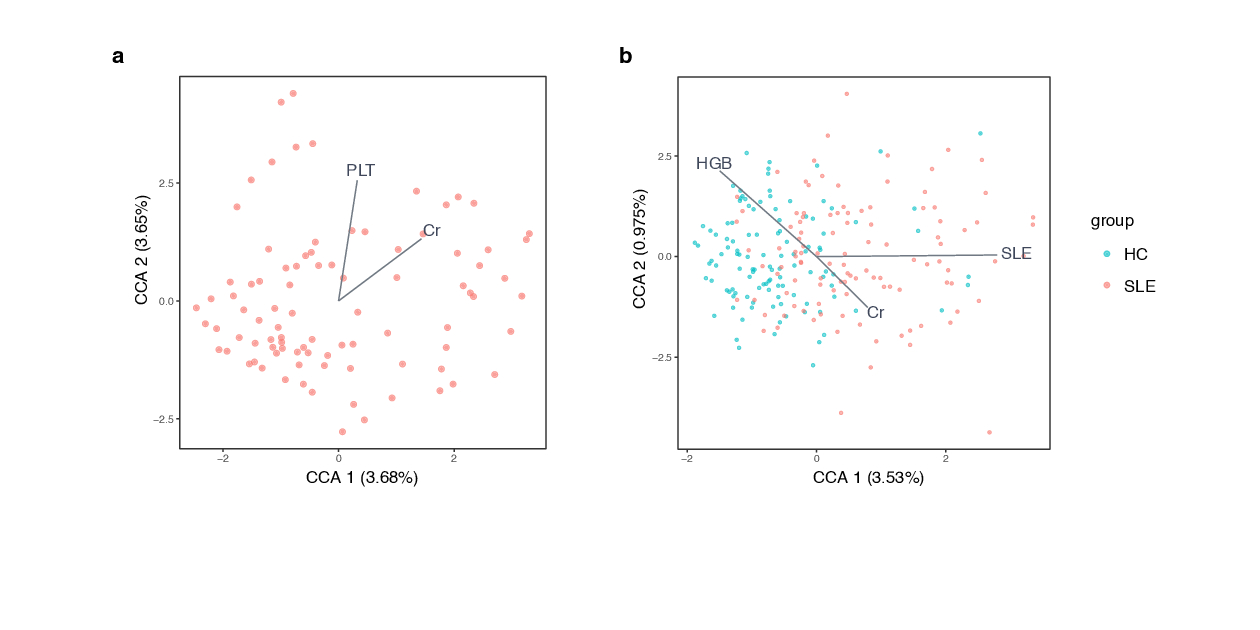

**Supplementary Figure 2. Constrained ordination analyses of the fecal microbiota of systemic lupus erythematosus (SLE). a,b,** Distance-based redundancy analysis (dbRDA), based on Bray-Curtis similarity, of the fecal microbiota of **(a)** 89 treatment-naive SLE patients or **(b)** 115 healthy control and 110 SLE patients with complete clinical data. Statistical significance was calculated by permutational multivariate analysis of variance (PERMANOVA) with a permutation number of 999. Clinical variables included, along with their effect size and significance, can be found in Supplementary Table 3. Plots show clinical variables that can substantially explain the variation between fecal samples.

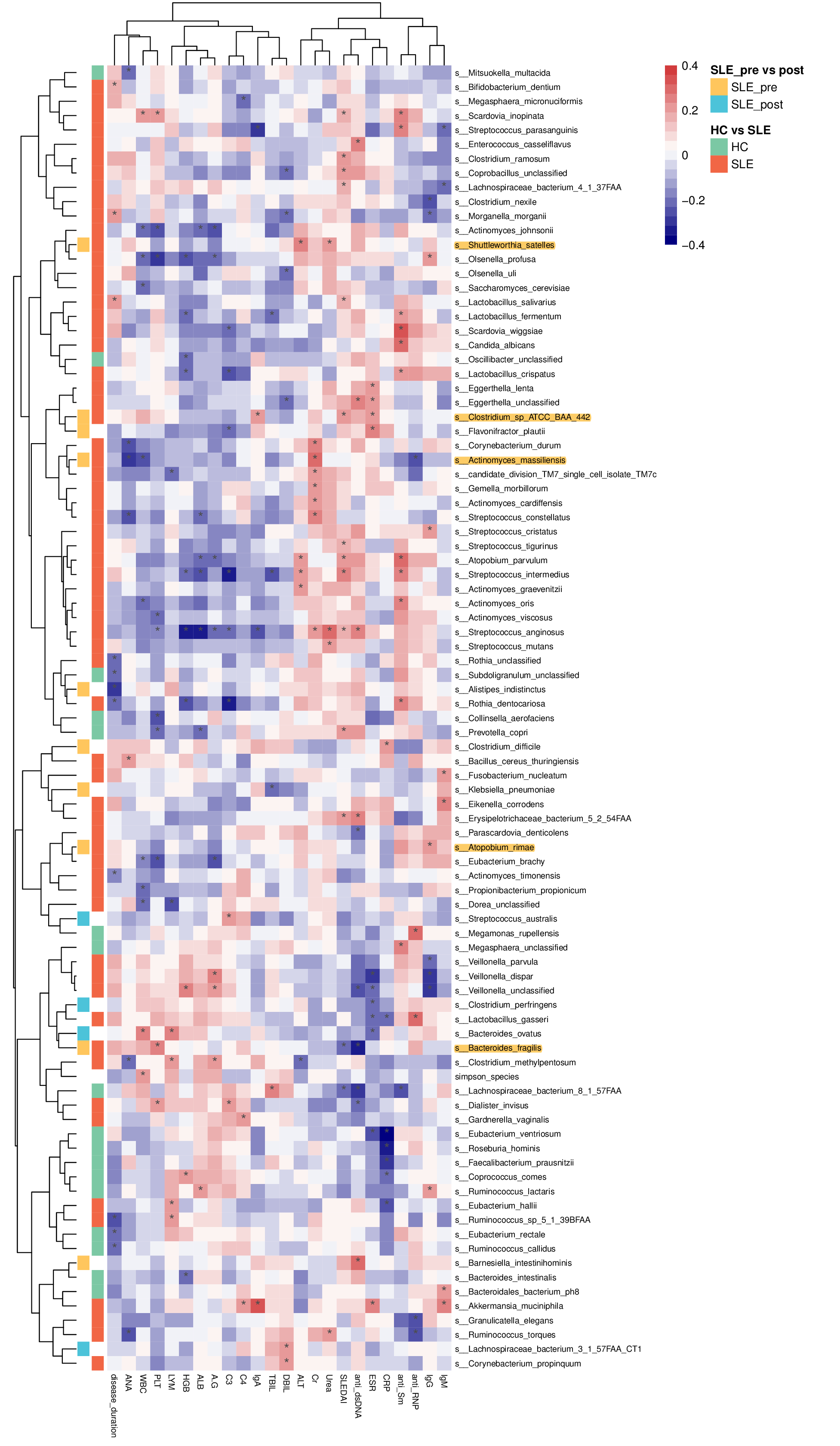

**Supplementary Figure 3. Correlation between taxonomical abundance and clinical data in patients of systemic lupus erythematosus (SLE).** Heatmaps of Spearman correlation between relative abundance of the gut species and clinical metadata. Taxa that were differentially enriched between SLE patients and healthy controls, or between pre- and post-treatment SLE patients, were included in the correlation analysis. Taxa that were not significantly associated with any clinical index were excluded from the heatmaps. Each row represents a different taxon while each column represents a different clinical index. Cell colors in the main heatmaps correspond to the value of correlation coefficients, with firebrick red represents a positive and navy blue represents a negative correlation. Asterisk shows a significant correlation (P＜0.05). Colors of cells in the left panel depict which group a specific taxon was enriched in. Green cells for healthy controls (HC); orange cells for all treatment-naive SLE patients; yellow cells for the pre-treatment SLE patients; blue cells for the post-treatment SLE patients. WBC, white blood cell; LYM, lymphocyte; HGB, hemoglobulin; PLT, platelet; ALT, alanine aminotransferase; ALB, albumin; TBIL, total bile acid; DBIL, direct bile acid; Cr, creatinine; CRP, C-reactive protein; ESR, erythrocyte sedimentation rate; ANA, antinuclear antibody; SLEDAI, SLE disease activity index.

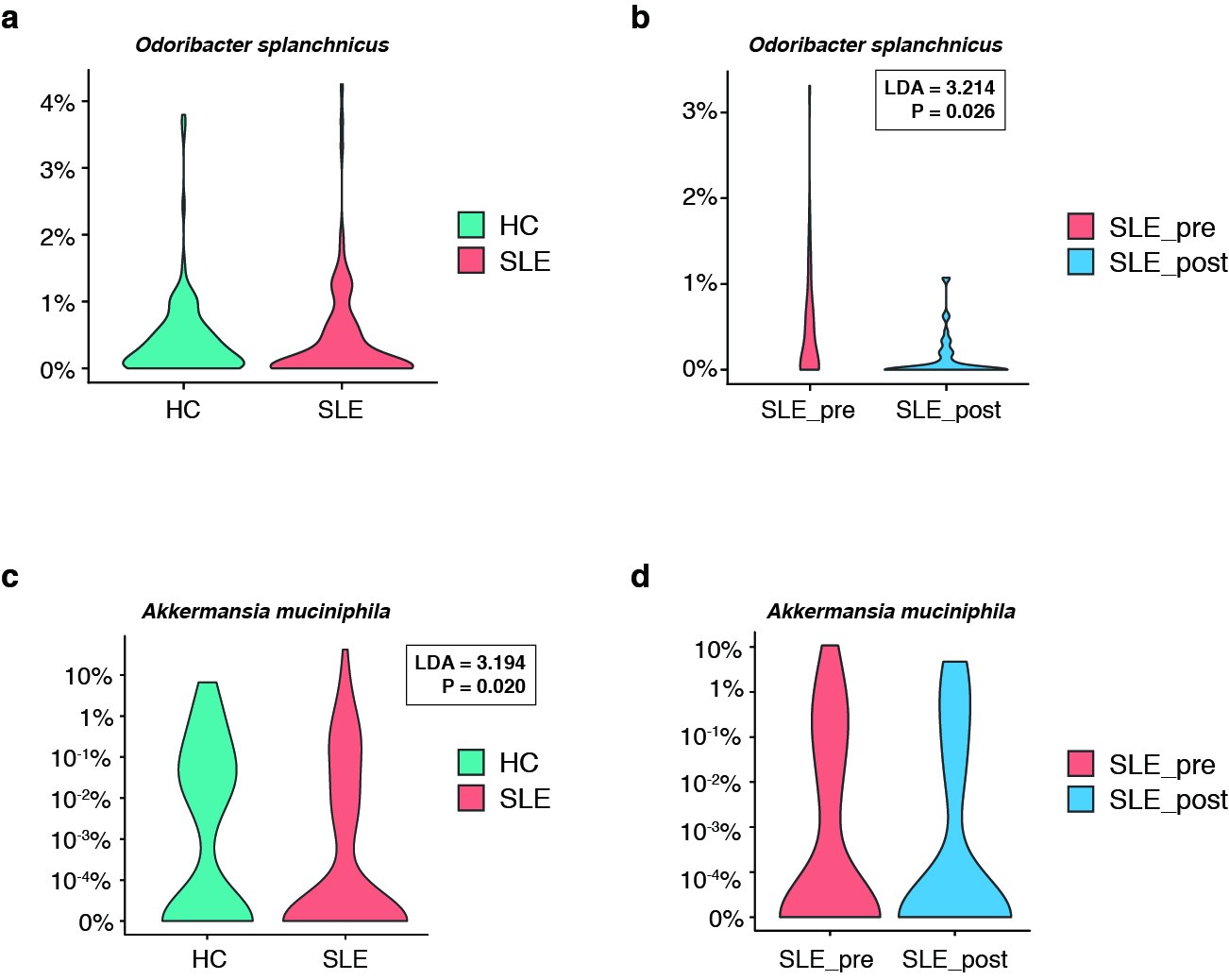

**Supplementary Figure 4. The relative abundance of** **gut *Odoribacter splanchnicus* between groups. a,b,** Violin plots showing the different relative abundance of *Odoribacter splanchnicus* **(a)** between the fecal samples of 115 healthy controls and 117 treatment-naïve systemic lupus erythematosus (SLE) patients (not significant), and **(b)** between 52 pre-treatment and post-treatment fecal samples of SLE patients (P = 0.026). **c,d**, Violin plots showing the different relative abundance of *Akkermansia muciniphila* **(c)** between the fecal samples of 115 healthy controls and 117 treatment-naïve systemic lupus erythematosus (SLE) patients (P = 0.020), and **(d)** between 52 pre-treatment and post-treatment fecal samples of SLE patients (not significant).The width of the violins reveals the data density at different abundance. The statistical significance and effect size were decided by linear discriminant analysis (LDA) of effect size (LEfSe).
